# Does PERM1 Regulate Systolic Cardiac Function? A Game of Numbers

**DOI:** 10.1101/2024.10.08.617184

**Authors:** Alexey V. Zaitsev, Karthi Sreedevi, Brianna Goode, Junco S. Warren

**Affiliations:** Fralin Biomedical Research Institute at Virginia Tech Carilion, Virginia Tech, Roanoke, VA, USA; Center for Vascular and Heart Research, Fralin Biomedical Research Institute, Virginia Tech, Roanoke, VA, USA; Department of Human Nutrition, Food and Exercise, Virginia Tech, Blacksburg, VA, USA

**Author notes:** Corresponding author: Junco S. Warren, Ph.D, **Email:**.

## Abstract

We and others demonstrated that PERM1 is a positive regulator of mitochondrial bioenergetics in the heart. However, discrepant results have emerged with regard to whether PERM1 loss-of-function affect cardiac contractility. In order to exclude the possibility that the reported negative results can be due to insufficient power of statistical test, we conducted a more robust echocardiography (Echo) analysis by increasing the sample size. We used *Perm1*-KO and their respective wildtype (WT) littermates, which were destined to tissue harvest. This yielded 84 WT mice and 88 *Perm1*-KO mice. We analyzed Echo-derived parameters of left ventricular (LV) systolic function. At the end of the study, ejection fraction (EF) was 65.43 ± 7.13 in WT vs. 53.98 ± 8.80 in P*erm1*-KO yielding p < 0.00000000000000004. Other parameters which reached statistically significant difference between WT and *Perm1*-KO (at p < 0.05) included LV fractional shortening (FS), LV diastolic and systolic diameters, LV anterior and posterior systolic wall thickness, LV posterior wall systolic thickening, stroke volume, and cardiac output (CO). Retrospectively, a p value < 0.05 was consistently achieved in assessment of EF only after average N per group reached 13. Larger minimal N per group were required for other parameters. Of interest, in both groups there were no correlation between EF% and CO. At the same time, in both groups EF strongly inversely correlated with LV diastolic diameter. This led us to a speculation that low EF may be in part compensated by an increased LV circumference, for the purpose of maintaining invariant CO. Indeed, the intergroup difference in CO (6%) was much smaller than the intergroup difference in EF (18%). We conclude that PERM1 does regulate cardiac mechanics. Changes caused by constitutive *Perm1*-KO can be conceptualized as reduced contractility partially compensated by increased LV circumference. This study underscores the importance of sufficiently large sample size for detecting significant differences in Echo data.

## Introduction

PERM1 (*<u>P</u>GC-1 and <u>E</u>RR regulator in <u>m</u>uscle <u>1</u>*) was discovered by Krali’s group [1] as a regulator of energy metabolism in skeletal muscle. Subsequent studies by our group and others demonstrated that this protein plays similar roles in energy metabolism of cardiac muscle [2–6]. We demonstrated that PERM1 is downregulated in both human and mouse hearts with significantly reduced systolic function [2]. However, whether downregulation of PERM1 contributes to reduced contractility in the failing heart remains unclear.

Current studies show conflicting results regarding the effect of PERM1 loss-of-function in mice and its role in regulating cardiac contractility. Whereas we previously reported that constitutive Perm1 deletion causes a decrease in cardiac systolic function [3], Huang et al. using exactly the same mouse model reported the lack of such effect [6]. Given that PERM1 is emerging as a potential therapeutic target for management of heart failure, it is important to determine unequivocally if, besides its effects on cardiac energy metabolism, PERM1 can also modulate cardiac mechanics.

The recent relocation of our group to the Fralin Biomedical Research Institute (FBRI) at Virginia Tech in Roanoke, VA required re-establishment of the entire framework and baseline for comprehensive analysis of cardiac function. As a part of this effort, we decided to reconfirm echocardiographic results obtained previously in University of Utah [3]. At the new facility we had an advantage of unlimited and no-cost access to echocardiographic system which enabled us to perform assessments of systolic function in all *Perm1*- KO mice and their WT littermates destined to tissue harvest for all molecular biological purposes performed between April of 2022 and September of 2023. As a result, we achieved a possibly unprecedented sample size per experimental group (over 80) among the relevant published literature.

We believe that the outcomes of this study merit publication for three reasons. First, it confirms with brutal force our previous result [3] regarding involvement of Perm1 in regulation of cardiac function, by yielding p value of less than 0.00000000000000004 in the effect of *Perm1*-KO with regard to reduction in EF (and at a lesser statistical significance, other parameters of systolic function). Second, it reveals (possibly due to relatively large number of observations) hitherto unrecognized reciprocal relationship between LV circumference (a largely structural parameter) and ejection fraction (a largely functional parameter) which may be important for the system-level understanding of normal and abnormal cardiac contractility. Lastly, it serves as a good reference for expected variability of outcomes in echocardiographic measurements for planning sample sizes in future studies.

## Methods

All animal experiments were approved by the Institutional Animal Care and Use Committee (protocol #21- 127) at Fralin Biomedical Research Institute at Virginia Tech.

*Perm1*-KO mice were generated by Crispr/Cas9 system with gRNA targeting at the exon 2 in the Sadoshima lab at Rutgers New Jersey Medical School, as previously described [3, 6]. This study included essentially all *Perm1*-KO mice (KO) and their wildtype littermates (WT) raised in our mouse colony at Virginia Tech and sacrificed between April of 2022 and September of 2023 for collecting a pool of tissue samples. Both male and female mice were included (n=39 in male WT, n=45 in female WT, n=40 in male KO, n=48 in female KO). We followed the guidelines established in a recently published consensus statement on echocardiography for assessments of systolic function in rodents [7]. All included Echo measurements were obtained in mice meeting the following criteria: heart rate between 450 and 560 beats per minute; core body temperature between 36 and 38 degrees Celsius; respiratory rate between 30 and 120 per minute. Mice were induced at 4% isoflurane in pure oxygen and were maintained at 1.5 – 2.5%, with slight adjustments to ensure that the vital parameters are within the required range. The mice were placed on a heated temperature-controlled platform and additional heat was provided from an infrared heating lamp with adjustable power. Measurements were obtained using Visual Sonics Vevo-2100 system with MS400 (38Mz) transducer. The measurements presented in this study were obtained in PSAX orientation using M-mode and analysis was performed using Visual Sonics Vevo LAB software ver. 5.7.0. While the number of Echo modalities and the total duration of measurements varied over the course of this study, the measurement presented here were obtained between 10 and 30 minutes after the onset of anesthesia. All the parameters presented int this study were retrieved from reports produced by Visual Sonics Vevo LAB software, with the exception of left ventricular (LV) posterior wall (LVPW) thickening, which was calculated as the difference between LVPW thickness in systole and diastole presented as the percentage of LVPW thickness in diastole. Unpaired two-sided t-test and linear regression analysis were performed using Data Analysis ToolPac in Microsoft Excel software. The following test for normality of distribution were performed using GraphPad ver. 10.2.3 (GraphPad Software LLC, Boston, MA): D’Agostino & Pearson test, Anderson-Darling test, Shapiro-Wilk test, and Kolmogorov-Smirnov test. For all measured Echo parameters both unpaired t-test and non-parametric Mann-Whitney test were applied to test for significance of difference between WT and KO groups. p-values less than 0.05 were considered statistically significant. Standardized mean difference (difference between means divided by common within-population standard deviation) was used as an estimator of effect size.

Whereas in general the endocardial boundaries of the LVAW and LVPW are relatively sharp in M-mode Echo images, and hence the estimates of LV diameter and volume can be treated with a relatively high degree of confidence, the epicardial boundaries of LV wall are often fuzzy and much less defined which can introduce a larger measurement error in estimates of LV wall thickness. Because of this, we surmised that a selection of a subset of data in which the epicardial LV boundaries are sharp and induce subjective confidence may achieve a better contrast in LV thickness estimates despite the reduction of the sample size. To test this conjecture, a researcher blinded to the mouse genotype (B.G.) was tasked with selection of mice in which either the epicardial LVAW boundary, or the epicardial LVPW boundary or both, were subjectively sharp and unequivocal. The results from the subsample obtained in such manner were compared to those in the full sample.

## Results

### Perm1 deletion causes remodeling of cardiac geometry and function

Table 1 presents the results from total of 84 WT and 88 *Perm1*-KO mice (the full set of data can be found in **S1 Spreadsheet**). Among the measured parameters, only heart rate, age, and diastolic thickness of LV anterior and posterior walls were not significantly different between *Perm1*-KO and WT mice. All other parameters were significantly different, with the probability of intergroup difference due to chance ranging from a fraction of a percent (in case of stroke volume and CO) to less than one in quadrillion (in case of EF and FS). Percentwise, the largest differences were observed in LV systolic volume (56.1 %), LV systolic diameter (19.5 %), and FS (22.0 %). The smallest differences among statistically significant differences were observed in stroke volume (5.7%) and CO (5.9%).

**Table 1.** Statistical summary of all parameters of systolic cardiac function.

| **Parameter** | **WT** | **KO** | **%**<br>**difference** | **Both WT**<br>**and KO**<br>**normal**<br>**distribution?** | **p-value**<br>**by**<br>**Student t-**<br>**test** | **p-value by**<br>**Mann-Whitney**<br>**test** |
| --- | --- | --- | --- | --- | --- | --- |
| Heart rate<br>(bpm) | 514.5 ± 33.5 | 512.3 ± 30.1 | 0.4 | No | NS | NS |
| Age (weeks) | 10.16 ± 1.03 | 10.8 ± 0.9 | 0.9 | No | NS | NS |
| LV diameter<br>systole<br>(mm) | 2.43 ± 0.35 | 2.90 ± 0.47 | 19.5 | Yes | 4.35E-12 | <0.0001 |
| LV diameter<br>diastole<br>(mm) | 3.76 ± 0.29 | 4.00 ± 0.37 | 6.4 | No | 5.05E-06 | <0.0001 |
| LV volume<br>systole (uL) | 21.60 ± 7.40 | 33.73 ± 13.89 | 56.1 | No | 3.29E-11 | <0.0001 |
| LV volume<br>diastole (uL) | 60.91 ± 11.14 | 70.79 ± 16.16 | 16.2 | No | 6.69E-06 | <0.0001 |
| Stroke<br>volume (uL) | 39.31 ± 5.60 | 37.07 ± 4.42 | 5.7 | No | 3.91E-03 | 0.0081 |
| Ejection<br>fraction (%) | 65.43 ± 7.13 | 53.98 ± 8.81 | 17.5 | Yes | 5.21E-17 | <0.0001 |
| Fractional<br>shortening<br>(%) | 35.58 ± 5.36 | 27.77 ± 5.55 | 22 | Yes | 4.11E-17 | <0.0001 |
| Cardiac<br>output<br>(mL/min) | 20.19 ± 2.89 | 19.00 ± 2.60 | 5.9 | Yes | 4.95E-03 | 0.0082 |
| LVAW<br>thickness<br>systole<br>(mm) | $1.35 \pm 0.24$ | $1.21 \pm 0.19$ | 10.3 | Yes | 4.97E-05 | 0.0002 |
| LVAW<br>thickness<br>diastole<br>(mm) | $0.90 \pm 0.16$ | $0.85 \pm 0.14$ | 4.8 | Yes | NS | NS |
| LVPW<br>thickness<br>systole<br>(mm) | $1.20 \pm 0.23$ | $1.08 \pm 0.17$ | 10.3 | Yes | 6.58E-05 | 0.0002 |
| LVPW<br>thickness<br>diastole<br>(mm) | $0.77 \pm 0.16$ | $0.74 \pm 0.14$ | 3.4 | No | NS | NS |
| LVPW<br>thickening<br>(%) | $1.58 \pm 0.15$ | $1.47 \pm 0.16$ | 6.9 | Yes | 1.07E-05 | <0.0001 |

The relatively large sample size used in this study gave a rare opportunity to apply tests for normality of distribution, and build frequency distribution histograms, which is hardly possible for typical sample sizes found in relevant literature (n < 20). We considered those cases in which all four normality tests failed to find a significant deviation from normal distribution in both WT and KO groups as a good assurance that the underlying distribution is normal in both experimental groups and hence application of t-test is appropriate. The Echo parameters with met this criterion include LV diameter in systole, EF, FS, cardiac output, LVAW thickness in systole, LVAW thickness in diastole, LVPW thickness in systole, and LVPW thickening. The parameters which did not meet this criterion include LV diameter in diastole, LV volume in systole, LV volume in diastole, stroke volume and LVPW thickness in diastole. We did not analyze the issue of normality in further detail, but importantly application of parametric t-test and non-parametric Mann-Whitney test yielded matching outcomes in regard to inference of statistically significant difference (see Table 1).

Fig. 1 displays distribution histograms for the three selected parameters EF, CO, and LVPW thickening. EF is the most often reported parameter of the systolic function, and also a parameter used clinically as a criterion for the diagnosis of heart failure with reduced ejection fraction [8]. CO, is, literally and figuratively, the net result of cardiac function as a service to the entire body. LVPW thickening seems to be a reasonable way to estimate the magnitude of ventricular wall deformation which reflects the active properties of cardiac muscle.

**Fig. 1.**
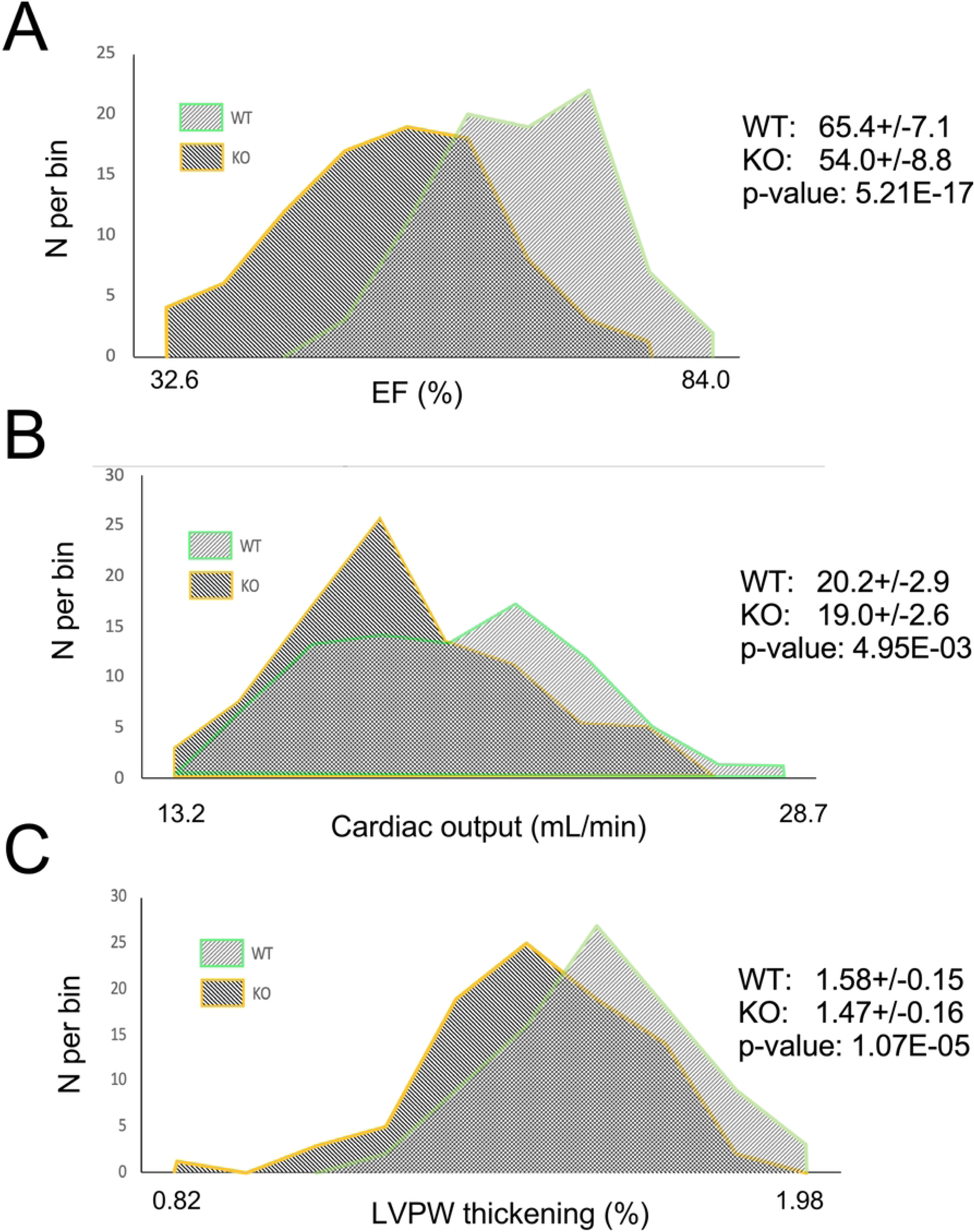
Frequency distribution histograms for selected parameters of systolic function. Frequency distribution histograms for EF (**A**), CO (**B**) and LVPW thickening (**C**) obtained in WT and *Perm1*- KO mice. Note that all distributions are unimodal and roughly bell-shaped. There is a big overlap in all distributions between WT and KO, yet the differences between means are statistically significant.

One can see that all 6 distributions are unimodal and more (as in case of EF) or less (as in case of CO and LVPW thickening) bell-shaped. The shapes of all 6 distributions are not significantly different from normal distribution according to all four normality tests (see Table 1). Hence, using the parametric Student’s t-test is appropriate. Despite large overlap between WT and KO in the distributions of all 3 parameters, the difference between means is statistically significant for all of them. It is of interest that all of WT mice have EF above 50%, which corresponds to the normal range in humans [1], whereas about 7% of *Perm1*-KO mice have EF below 40%, which corresponds to the standard definition of heart failure with reduced EF (HFrEF) [8]. Do these mice have cardiac dysfunction? Considering that cardiac output is the ultimate measure of cardiac performance, we tested whether low EF translates into low CO. Figure 2 shows that this is not the case. Fig. 2A plots CO as a function of EF in WT and KO mice. One can see that there is no correlation between these two parameters neither in WT nor in *Perm1*-KO mice. Essentially the same range of cardiac outputs is observed at EF < 40% and at EF > 70%. Even though the difference in CO between WT and KO does reach statistical significance, the magnitude of this difference (∼6 %) is much less than the magnitude of the difference in EF (∼18%). In this regard, it is interesting to note that in both WT and KO groups there is a strong inverse correlation between LV diastolic diameter and EF (Fig. 2B). LV diastolic diameter can be considered as a parameter mostly reflecting structural (passive) myocardial properties, whereas EF is widely perceived as the parameter mostly reflecting functional (active) myocardial properties. The data shown in Fig. 2B suggests that in both WT and *Perm1*-KO mice there is a reciprocal relationship between passive and active properties such that lower contractility is somewhat compensated by a larger lumen of LV chamber.

**Figure 2.**
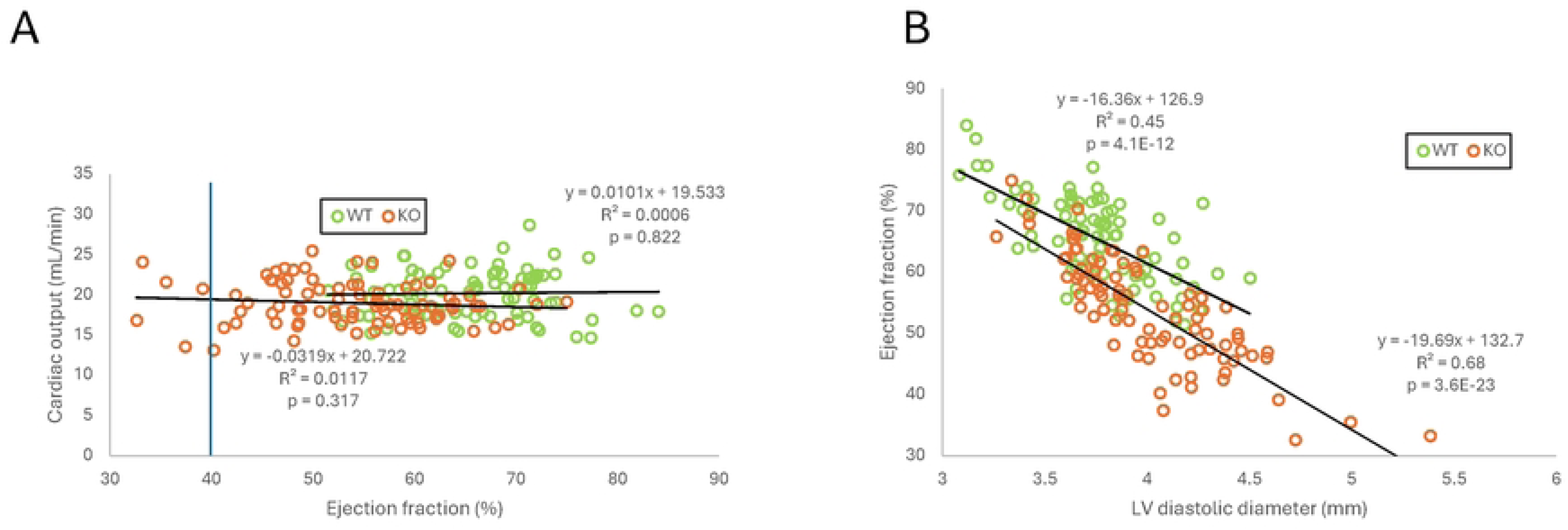
The analysis of correlation between EF and CO and between LV diameter in diastole and EF. Perm1 deletion does not change the relationship between EF and CO (**A**) and between LV diameter in diastole and EF (**B**). In both WT and *Perm1*-KO mice, there is no correlation between EF and CO (**A**), and there is a strong negative correlation between LV diameter in diastole and EF (**B**).

Fig. 3 presents the raw PSAX M-mode recordings from 4 mice having the lowest and the largest EF in WT mice (Figure 3, A and B respectively) and having the lowest and the largest EF in KO mice (Fig. 3C and 3D respectively). These images help to visualize the notion that in both WT and *Perm1*-KO mice hearts with relatively “dilated” LV and low EF effectively achieve the same stroke volume and cardiac output as hearts with more compact LV and higher EF. This relationship appears to be similar in WT and KO groups; however, the distribution of LV diastolic diameter is shifted to higher values, and the distribution of EF is shifted towards lower values in KO as compared to WT.

**Figure 3.**
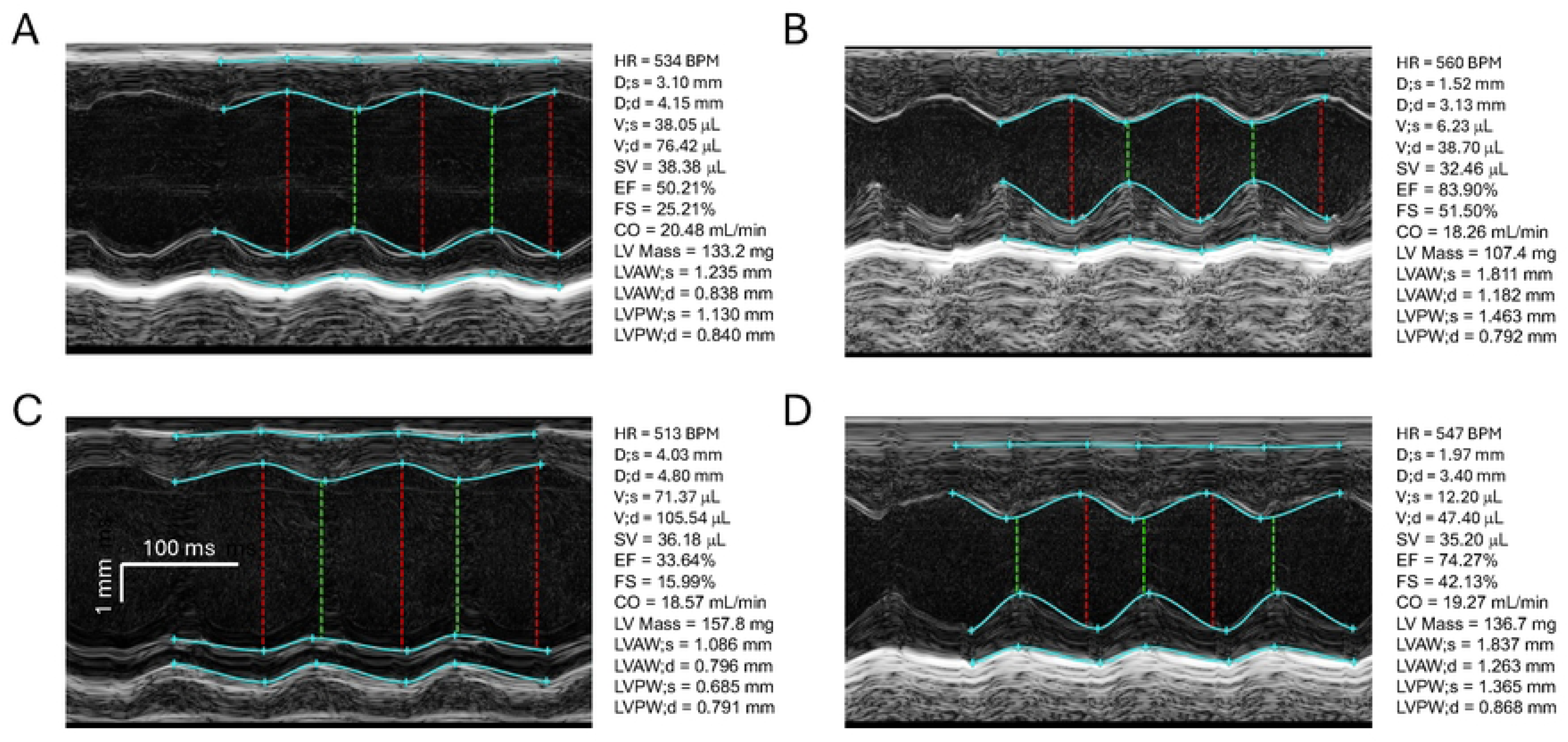
Echocardiographic images representing the extremes in the distribution of EF in WT and *Prem1*-KO groups. PSAX M-mode images representing the mice with the lowest (**A**) and the highest (**B**) value of EF in the WT group, and the lowest (**C**) and the highest (**D**) value of EF in the KO group. Note that despite the very big differences in EF values, these mice, representing the extreme ends of EF distributions, achieve similar levels of CO.

### Statistical considerations in analysis of differences in systolic function between WT and Perm1-KO

The unprecedently large sample size achieved in this study may be useful for power analysis necessary (and required by funding agencies) to plan numbers of animals per group to be used in future studies. In power analysis, researchers consider the effect size and the acceptable levels of Type I error (false positives) and Type II error (false negatives). The effect size is essentially the relationship between the differences in mean values and the dispersion of outcomes in experimental groups. Here we use a qualitative categorization of the effect size proposed and discussed in [9]. Table 2 presents the estimates of effect size and the minimal N per group assuming the accepted probability of Type I error (alpha) of 0.05 and three different levels of probability of Type II error (beta): 0.2, 0.1, and 0.05. It is interesting to note that the most widely used acceptable values for alpha and beta are 0.05 and 0.2 respectively. In essence, researchers typically allow for much larger chance of missing a significant difference than mistakenly accepting a difference which does not exist. This is purely a matter of choice, and we owe this specific choice largely to an influential publication of Cohen et al. [10] One could argue that the acceptable chance of false negative should be the same as the acceptable chance of false positive. But again, this is entirely up to the researcher. Based on the data collected in this study, Table 2 provides minimally necessary sample sizes for different choices of beta (at the fixed level of alpha = 0.05, which is so deeply ingrained in modern culture of biomedical research that we will accept it as a strict constant).

**Table 2.** Sample size necessary for detection of significant differences in specific parameters of systolic function at different levels of type II error (beta.

| Parameter | % diff. | Effect size | Effect size category | Required N per group (alpha = 0.05) |  |  |
| --- | --- | --- | --- | --- | --- | --- |
|  |  |  |  | beta = 0.2 | beta = 0.1 | beta = 0.05 |
| LV diameter<br>systole (mm) | 19.5 | 1.13 | large | 14 | 18 | 22 |
| LV diameter<br>diastole (mm) | 6.4 | 0.72 | medium | 32 | 42 | 51 |
| LV volume<br>systole (uL) | 56.1 | 1.09 | large | 15 | 19 | 23 |
| LV volume<br>diastole (uL) | 16.2 | 0.71 | medium | 32 | 43 | 53 |
| Stroke volume<br>(uL) | 5.7 | 0.44 | small | 81 | 108 | 133 |
| Ejection<br>fraction (%) | 17.5 | 1.43 | very large | 9 | 12 | 14 |
| Fractional<br>shortening (%) | 22.0 | 1.43 | very large | 9 | 12 | 14 |
| Cardiac output<br>(mL/min) | 5.9 | 0.43 | small | 85 | 114 | 140 |
| LVAW<br>thickness<br>systole (mm) | 10.3 | 0.65 | medium | 39 | 52 | 64 |
| LVPW<br>thickness<br>systole (mm) | 10.3 | 0.59 | medium | 46 | 61 | 75 |
| LVPW<br>thickening (%) | 6.9 | 0.71 | medium | 33 | 43 | 53 |

Of interest, the sample size per group achieved in this study is actually close to the size minimally sufficient to detect significant differences in *all* listed parameters at the level of alpha = 0.05 and the level of beta = 0.2. At a typically reported sample size of 15 - 20 per group, and accepting alpha = 0.05 and beta = 0.2, we could hope to detect significant differences in only four out of measured parameters: LV systolic diameter, LV systolic volume, EF and FS. Fig. 4 shows the historical analysis in which the p-value yielded by unpaired two-sided t-test is plotted against the average group size between WT and KO groups, as the group sizes grew over time. These data are plotted for the same parameters as shown in Fig. 1: EF, CO, and LVPW thickening. It is of interest that negative logarithm of p-value grew linearly with the sample size. For each analyzed parameter, the p-value fluctuated as the sample size grew, but after reaching a certain sample size, the p-value never returned below the value of 0.05. For the EF, the minimal average group size to achieve an assurance of p < 0.05 was n = 12.5, and for both CO and LVPW thickening it was n = 30.5

**Figure 4.**
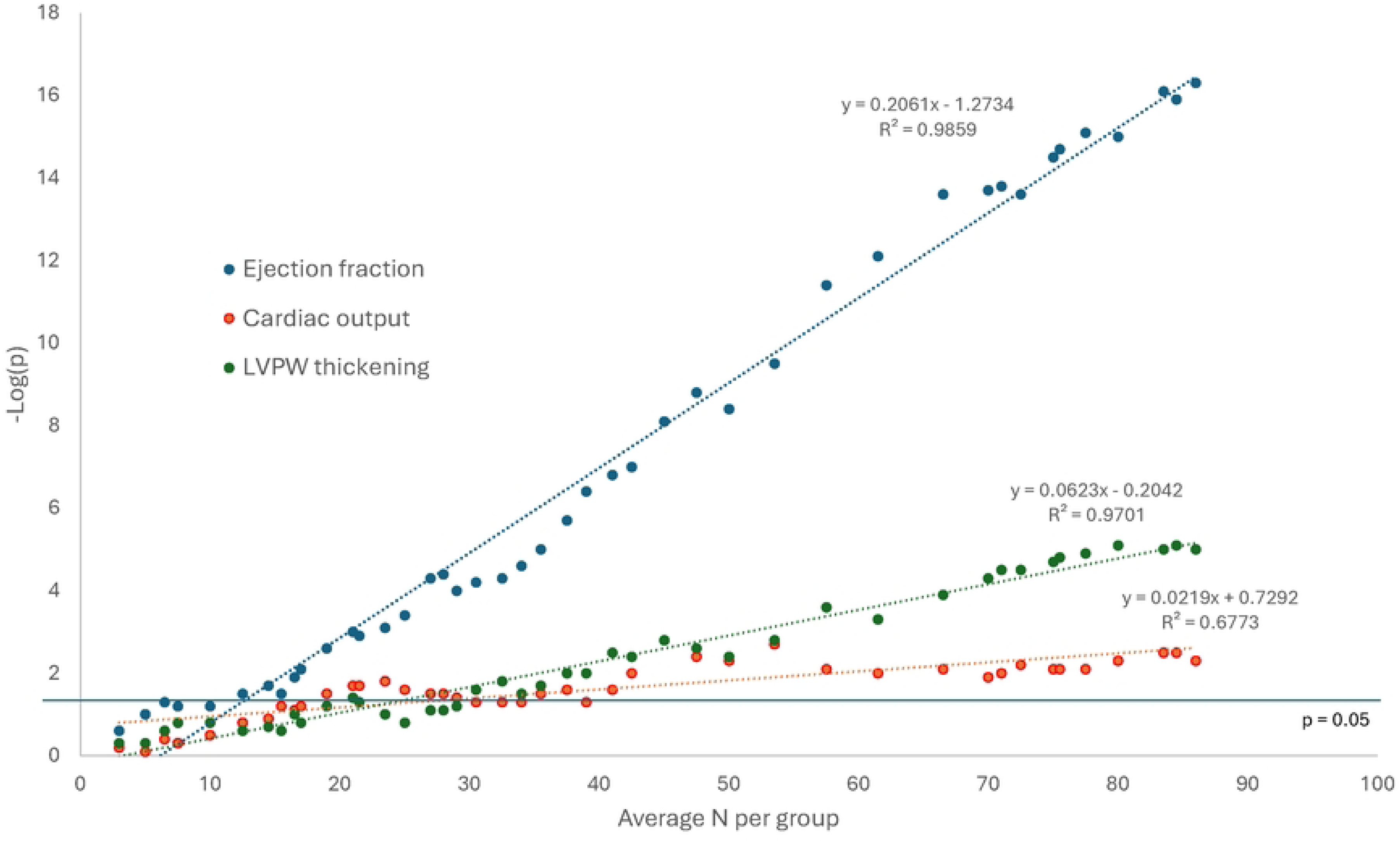
Retrospective analysis of p-value versus average group size. The p-value for unpaired two-sided t-test was computed as if measured each time the sample size increased due to additional experiments. The value of negative logarithm of p value increases approximately linearly with the average group size, and after group size reaches a critical number, the p-value remains at the level < 0.05.

Selection of cases with “good” LVAW epicardial boundary and “good” LVPW epicardial boundary did not reveal a significant difference in any parameter which would not be statistically significant in the full sample, but lost statistical significance in estimation of stroke volume, CO and LVPW thickening, presumably due to reduced sample size and hence the power of the statistical test (data not shown). From this we conclude that the “curation” of Echo data based on perceived sharpness of wall boundaries perhaps cannot help in increasing the sensitivity of the analysis at the expense of sample size.

## Discussion

The most important result of this study is incontrovertible evidence that *Perm1* deletion substantially affects cardiac geometry and function. The percent difference between the control and test groups ranged from approximately 6% for CO, stroke volume, LV diastolic diameter, and LVPW thickening to over 50% for LV systolic volume. In the middle of this range, EF, FS, LV systolic diameter and LV diastolic volume stand at about 20% difference between WT and *Perm1*-KO groups. Are these differences big or small? From a clinical perspective, EF is by far the most studied and reported parameter of cardiac systolic function in the clinical setting. Several studies reported that differences in EF of 5 percent units or more are clinically significant as predictors of morbidity and mortality in heart failure patients [11,12]. Hence, the absolute difference of > 10 percent units observed in this study should be considered clinically significant. Other parameters of systolic function reported here are rarely analyzed in the clinical setting. In one study the authors analyzed stroke volume in patients with aortic stenosis and concluded that a reduction 70 ml to 55 ml (∼20% change from healthy controls) was associated with a significantly increased risk of mortality, whereas the lesser degree of reduction was not [13]. In this setting, a reduction of stroke volume of ∼6% observed in our study would probably not be clinically significant, with coincides with our intuitive assessment that such a decrease in stroke volume is probably of little physiological relevance. We also found a study which reported a 5% to 8% decrease in LV systolic diameter and LV diastolic diameter in patients with heart failure and a reduced EF (HFrEF) who were treated with sodium-glucose cotransporter-2 inhibitors (SGLT2i) [14]. The authors cautiously suggested that these modest changes in LV volumes could contribute to improved clinical outcomes following SGLT2i administration in this cohort [14]. Hence, the increases in LV systolic diameter (56%) and LV diastolic diameter (16%) observed in *Perm1*-KO as compared to WT have a potential clinical relevance.

While the approximately 20% shift in the cental location of EF distribution towards lower values resulted in some *Perm1*-KO mice having EF values below 40%, leading to a potential conclusion of “cardiac dysfunction”, this interpretation should be moderated by the fact that CO, a more definitive measure of cardiac performance, is only reduced by 6%. Importantly, *Perm1*-KO mice show no signs of overt pathology. Thus, it may be more appropriate to describe the changes induced by *Perm1* deletion as remodeling rather than dysfunction. This remodeling affects both the passive and the active properties of the left ventricle in a complex way. Although the remodeled hearts exhibit increased left ventricular diastolic volume and reduced EF, these changes partly offset each other, resulting in only a minor decrease in cardiac output. Nevertheless, our current study underscores the role of PERM1 in regulating systolic function, as further supported by our recent adeno-associated virus (AAV)-mediated gain-of-function study [15].

Cho et al. demonstrated that PERM1 expression increases in mouse embryonic and postnatal hearts, suggesting a role of PERM1 in heart maturation [5]. In constitutive *Perm1*-KO mice, it is possible that some changes occurred earlier in the heart maturation, and some were an adaptive response to the primary changes. Another possibility is that the primary effects of *Perm1* deletion affected the active properties of cardiac contraction, and the relative dilation of the LV was an adaptive response to reduced contractility in an attempt to maintain adequate cardiac output. However, the opposite sequence of events cannot be ruled out. Supporting the potential direct effect of PERM1 on cardiac contractility, our recent study showed that adeno-associated virus (AAV)-mediated overexpression of PERM1 enhances systolic function in mice, in association with PERM1’s interaction with Troponin C, a cardiac-specific contractile protein [15]. This positive effect on cardiac contractility was observed in healthy mouse hearts, where ATP production and utilization are not impaired. Therefore, both our PERM1 loss-of-function and gain-of-function studies consistently suggest a role for PERM1 in regulating cardiac contractility. This will need to be further confirmed in future studies using inducible, cardiac-specific *Perm1*-KO mice.

The results of this study, showing a subtle yet significant reduction in EF in *Perm1*-KO mice compared with their WT littermates, confirm our previously published results [3]. Notably, the average of EF in both WT and *Perm1-*KO mice was slightly lower in the results obtained at the University of Utah. This difference might be attributed to subacute or chronic effects of high altitude (∼1600 meters at the site of University of Utah) on heart function [16]. However, our current and previous studies partially diverge from the results reported by Huang et al. [6], who did not observe significant differences in several parameters of systolic function between *Perm1*-KO mice line #29 and their littermates, despite using the same mouse line as used in our study. The following parameters were analyzed in both their and our study: LV internal diameter in diastole, LV internal diameter in systole, LVPW in diastole, and EF. Among these parameters, both studies did not find significant difference in LVPW in diastole between WT and KO groups. However, unlike in the study by Huang et al., we found significant differences between WT mice and *Perm1*-KO mice in LV internal diameter in diastole, LV internal diameter in systole, and EF. Huang et al. used 21 WT and 14 *Perm1*-KO line #29 mice. Based on our power analysis (see Table 2), this smaller sample size (n = 14) could be a limiting factor in detecting differences in LV internal diameter in diastole and LV internal diameter in systole, as the probability of a false negative exceeds 20%. However, for EF, their sample size appears sufficient, as our estimates indicate a less than 5% probability of missing a significant difference in this parameter (Table 2). Therefore, the discrepancy in EF results between our studies is unlikely due to sample size limitations and may instead stem from methodological differences, such as the use of Avertin in Huang et al.’s study compared with isoflurane in ours, among other factors.

Any anesthetic, including isoflurane, is known to suppress cardiac contractility to some extent [17,18]. In order to alleviate this problem, some researchers perform Echo recordings in conscious mice, which are usually pre-trained and “gently” restrained in the necessary supine position [17]. Unfortunately, training can hardly prevent the enormous stress imposed on mice immobilized in the most vulnerable position imaginable for these prey animals. Hence, it cannot be excluded that the Echo results obtained in conscious mice are affected by stress [18]. One noticeable difference between EF data obtained in mice under isoflurane as compared to those obtained in conscious mice is a much smaller dispersion observed in the latter case (for example, compare Fig. 1M in [19] with Fig. 2A in [6]). So, the question arises whether larger dispersion recorded under isoflurane is an artifact of inter-animal sensitivity to the anesthetic, or whether the smaller dispersion recorded in conscious animals is an artifact of extreme stress driving cardiac contraction to the absolute physiological limit which would cause a “saturation” in measured values of EF. We are unaware of any systematic studies which would prove or refute any of these assumptions. We believe however that whereas differences and/or dispersion in EF could depend on the effects of anesthetic, it is less likely that the LV diameter in diastole which is mostly determined by structural, or passive, properties of the LV (essentially, the LV size) could be affected by anesthesia. Note that the difference between most compact heart from WT group (LV diastolic diameter = 3.13 mm, see Figure 3B) and the most “dilated” heart from KO group (LV diastolic diameter = 4.80 mm, see Figure 3C) is more than 50%. We are convinced that such difference cannot be explained either by an error of measurement or by the effect of anesthetic, and has to reflect the real difference in the anatomy of the LV in these mice.

## Acknowledgements

We thank Dr. Allison Tegge for her assistance with statistical analysis and valuable suggestions. We are also grateful to Dr. Robert Goudie for his generous support and for providing access to the echocardiography system in his laboratory. Additionally, we greatly appreciate the technical assistance provided by Jane L. Jourdan, Sydney Bui, and Katia Olmos.

## Supporting information

**S1 Spreadsheet.** All echocardiographic data.

